# Systems Immunology in Obesity: Uncovering Hub Gene Networks in Macrophage-Specific Drivers of Chronic Inflammation

**DOI:** 10.64898/2026.11.26.769525

**Authors:** Felipe Caixeta, Adriana Simizo, Helder Nakaya, Tatiani Uceli Maioli

## Abstract

Obesity is marked by chronic low-grade inflammation, with macrophages in adipose tissue acting as key mediators of immune dysfunction. Macrophages play a crucial role in this process, undergoing significant gene expression changes. Understanding the molecular mechanisms driving these changes is critical for developing effective therapeutic strategies, yet the transcriptional regulators driving this immunometabolic reprogramming remain poorly understood. To better understand the molecular drivers of macrophage dysregulation in obesity, we applied a systems biology approach focused on immune gene networks.

We combined bulk RNA-seq network analysis (WGCNA) with cross-species single-cell validation to identify conserved immune regulators in ATMs. Using murine and human scRNA-seq datasets (GEO), we mapped co-expression networks in obese versus lean conditions, prioritizing hub genes by betweenness centrality and participation coefficient.

Our analysis recovered known obesity-associated immune regulators like Stmn1, Cadm1, Asph, Myo1e, Sms, Pianp, Ndufs5, Ftl1, 180027I7Rik and uncovered four novel hub genes Gcsh, Rcan1, Ndufb4, and Ywhaq that were not previously linked to macrophage biology or obesity. These novel genes displayed strong network connectivity and were robustly expressed in macrophage populations across species, suggesting a conserved role in macrophage mediated inflammation. Our integrative approach underscores the utility of combining co-expression network analysis with cross-species single-cell validation to uncover novel immune regulators in metabolic disease contexts both in mice and human cells. The identification of novel hub genes, provides valuable insights into the molecular mechanisms underlying macrophage dysfunction in obesity and suggests potential therapeutic targets for combating this complex disease.

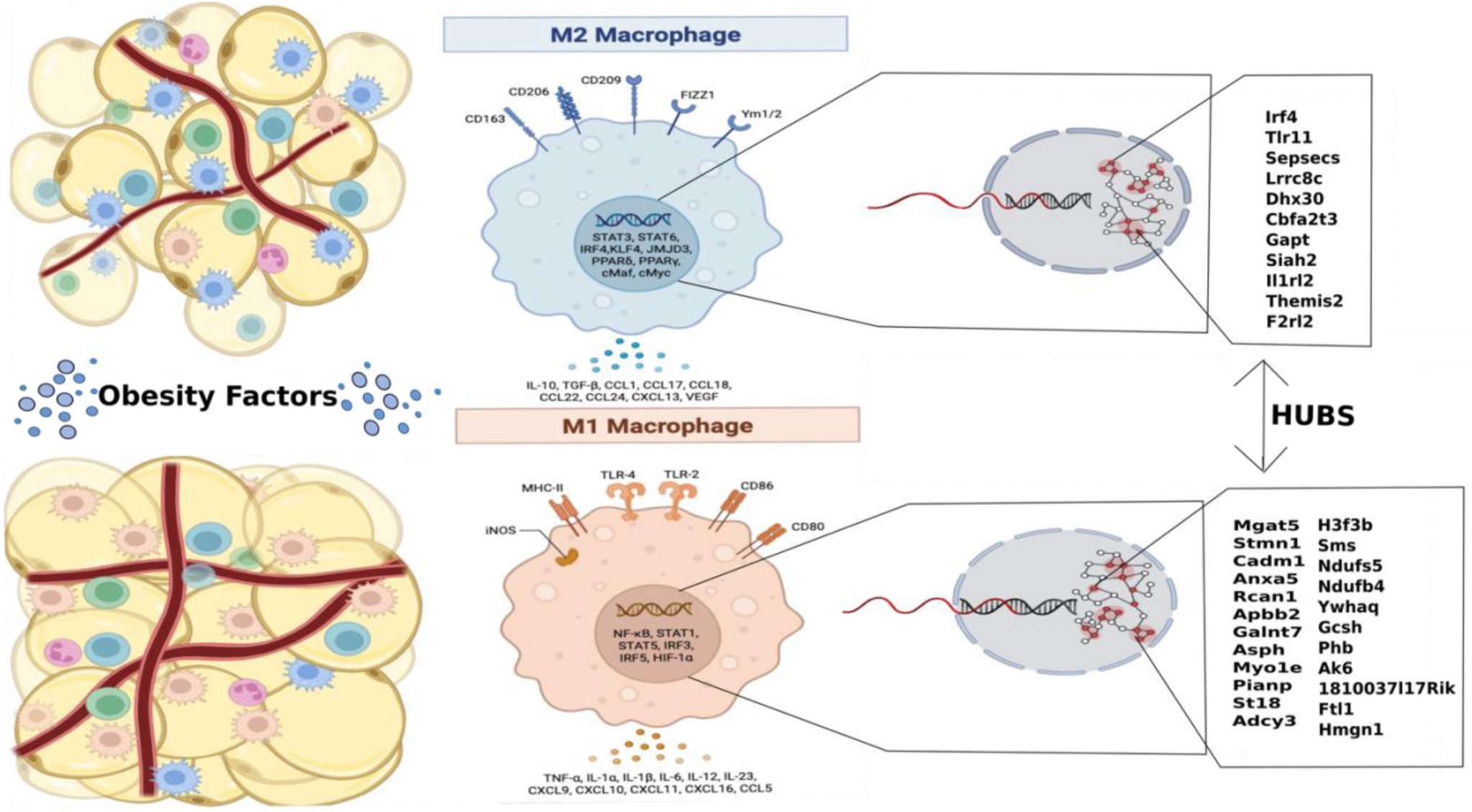

## Introduction

The evolutionary pressure from the historical scarcity of high calorie foods likely favored genetic pathways promoting energy intake and storage, predisposing modern humans to obesity. In the 21st century, the escalating global incidence of overweight and obesity has become a critical public health concern in most nations, yet effective treatment strategies remain limited^1^, mathematical models currently employed to analyze and predict disease prevalence have been applied to obesity, indicating that if current growth rates are maintained, all 185 countries studied will show an increase in obesity by 2030^2^.

Obesity is a clinically complex, multifactorial pathology defined by an excessive or anomalous distribution of adipose tissue, stemming from a fundamental dysregulation of appetite and energy metabolism, which compromises overall health^3^, as a complex multifactorial condition, obesity is associated with the occurrence of numerous comorbidities, including metabolic syndrome ^4^, cardiovascular diseases, and chronic inflammatory conditions such as type 2 diabetes mellitus^3^. It is also linked to non-alcoholic fatty liver disease^5^, hypertension^6^, dyslipidemia^7^, asthma^8^, both male and female infertility ^9,10^, and various types of cancer^11^. More recently, obesity has been identified as a significant risk factor for severe COVID-19^12^ and has been associated with increased susceptibility to infectious diseases in general^13^, consequently, obesity substantially reduces quality of life, shortens life expectancy, and elevates healthcare costs14. Due to its complex and chronic profile, obesity transcends a simple imbalance of energy intake and expenditure. It is recognized as a complex pathology involving metabolic alterations, oxidative stress, mitochondrial dysfunction, and immune dysregulation, which establish a state of low-grade chronic inflammation^15,16^.

The scientific understanding of adipose tissue has shifted significantly in the past three decades, paralleling the dramatic rise in the prevalence of obesity and its comorbidities, this tissue is capable of secreting a wide array of molecules, including leptin, adiponectin, cytokines such as TNF-α, IL-6, and IL-1β, and vascular regulators like angiotensin II and plasminogen activator inhibitor-1 (PAI-1)^17^. The expansion of adipose tissue during obesity is characterized not only by adipocyte hypertrophy but also by the infiltration of bone marrow-derived cells, with macrophages and neutrophils being among the first to arrive at the site^18^. The resident immune cells within adipose tissue, including T and B lymphocytes, dendritic cells, and particularly macrophages, become dysregulated in the context of obesity, contributing to the development of a perturbed immune system and the progression of the disease^19^.

Macrophages are pivotal to adipose tissue immunity, where they function as antigen-presenting cells and modulate T-cell responses^20^. In the lean state, resident adipose tissue macrophages (ATMs) comprise about 10% of immune cells, are distributed along the vasculature, and primarily exhibit an anti-inflammatory and tissue-remodeling phenotype^21,22^. A key function of these resident cells is the production of anti-inflammatory mediators such as IL-10, which supports the maintenance and expansion of regulatory T cells (Tregs)^19,23^. However, obesity induces a profound shift in the ATM population, number, and function, their population expands to represent up to 50% of the cells in adipose tissue, driven by the infiltration of circulating monocytes that aggregate in “crown-like structures” around dying adipocytes^21^. This infiltration leads to the predominance of a pro-inflammatory macrophage phenotype, characterized by elevated expression of genes for inflammatory mediators^22^. This phenotypic switch is a result of altered gene expression programs in response to the obese microenvironment^24^. Once activated, these macrophages secrete cytokines (e.g., TNF-α, IL-1β) and chemokines that amplify inflammation, recruit additional immune cells, and interact with other local lymphocytes and innate lymphoid cells (ILCs)^19^. This inflammatory signaling is regulated by intracellular cascades like the JAK pathway, which influences insulin sensitivity and is in turn modulated by suppressors of cytokine signaling (SOCS) proteins^25^, linking macrophage activation directly to the metabolic dysregulation seen in obesity. This phenotypic plasticity, leading to distinct macrophage populations that secrete different mediators, can be attributed to differential gene expression patterns induced by the obese microenvironment^24^.

With the proliferation of public gene expression data from high-throughput sequencing, a primary challenge is the extraction of biologically useful information. While differential gene expression (DGE) analysis is effective for identifying individual genes associated with a phenotype^22^, it does not account for the complex interactions between genes that drive cellular processes. Since biological functions arise from the coordinated action of multiple genes, a systems level understanding is necessary to elucidate the mechanisms underlying complex traits^26^. To overcome this limitation, systems level approaches such as network analysis are increasingly utilized^27–29^, gene co-expression network analysis provides such a systems level perspective. This approach operates on the principle that genes with correlated expression patterns are often functionally related^30^. These networks are widely used to infer gene function, identify novel regulatory pathways, and prioritize candidate genes. Although co-expression does not definitively prove co-regulation, its utility in functional genomics is well established and continues to grow with advancing sequencing technologies^31^. The application of this methodology is particularly pertinent to the study of obesity. It has been demonstrated that genes within a common metabolic pathway often share co-expression patterns^32^. As obesity is a complex metabolic disease characterized by chronic low-grade inflammation, there is strong evidence for an intrinsic crosstalk between metabolism and immunity. This relationship, however, remains incompletely understood. Gene co-expression analysis therefore offers a powerful framework to explore the transcriptomic landscape governing the interplay between metabolic and immune pathways in obesity.

In this study our goal was to address this gap, this study was conceived to leverage co-expression network analysis on publicly available transcriptomic data. The aim of this research is to construct a gene co-expression network from murine macrophages in an obese state to identify key genes that govern their functional and phenotypic polarization, providing novel insights into the regulatory mechanisms at play during obesity.

## Material and methods

### Data Acquisition for Gene Co-expression Network Construction

To construct the gene co-expression network, a search was performed in the NCBI Gene Expression Omnibus (GEO) database for studies containing transcriptome sequencing data of adipose tissue macrophages from obese and control mice. The dataset selected for this analysis was from Silva HM, et al. (2019)^33^, available under GEO accession number GSE126407. This study performed paired-end RNA sequencing (Illumina HiSeq 2500) on cDNA libraries prepared from adipose tissue of male C57BL/10.PL mice. In the source study, cellular populations (monocytes, macrophages, and dendritic cells) were isolated via flow cytometry and cell sorting. For the present analysis, only the data corresponding to the macrophage populations identified as VAM1, VAM2, and DPs were utilized.

### Murine Single-Cell RNA-seq Data for Hub Gene Analysis

To analyze the identified hub genes at the single-cell level, a publicly available murine single-cell RNA sequencing (scRNA-seq) dataset was obtained from GEO under accession number GSE117176 ^34^. This dataset contains transcriptomic profiles of adipose tissue macrophages isolated from mice fed either a control diet or an obesogenic high-fat diet (HFD). In the original study, macrophages were isolated using a BD Biosciences FACS Aria II cell sorter. The scRNA-seq libraries were prepared using the Chromium Single Cell 3’ Reagent Kit v2 (10x Genomics) on the GemCode Single Cell Instrument, and sequencing was performed on an Illumina HiSeq 4000 platform.

### Human Single-Cell RNA-seq Data for Comparative Analysis

For subsequent analysis in a human context, a scRNA-seq dataset of human adipose tissue macrophages was acquired from GEO under accession number GSE176171 ^35^. The macrophages were isolated from adipose tissue obtained from patients undergoing plastic surgery. Library preparation was conducted using the Chromium Single Cell 3’ v3.1 Chip (10x Genomics), and sequencing was performed on an Illumina NextSeq 500 instrument.

### RNA-seq Data Processing, filtering and analysis

All data processing and analysis were conducted within the R programming environment. The initial dataset (GEO accession: GSE126407) comprised 33 samples from various cell types. For this study, we selected only the samples corresponding to macrophage populations, specifically those annotated as double-positive (DPs), vasculature-associated macrophage 1 (VAM1), and vasculature-associated macrophage 2 (VAM2), resulting in a subset of 17 samples and 49,107 genes(Suplementary Figure 1A).

A multi-step quality control and filtering pipeline was then applied. First, sample integrity was assessed by checking for missing values and identifying outliers. A hierarchical clustering analysis revealed that one sample, VAM.2.ND.1(Suplementary Figure 1B), was a significant outlier and was consequently removed from the dataset (Suplementary Figure 1C). Next, a gene-level filtering was performed to remove genes with zero variance or no expression across all retained samples. After the removal of the outlier sample and non-expressed genes, the final dataset used for network construction consisted of 16 macrophage samples for lean and obese tissues and 29,767 expressed genes(Suplementary Figure 1D). Combining differential gene expression (DGE) analysis with co-expression network construction is a powerful strategy for elucidating the molecular mechanisms of biological processes. In this study, the DGE analysis was performed using the DESeq2 R package^36^ to identify individual genes with statistically significant changes in expression.

### Single-Cell RNA-seq Data Processing and Analysis

Analysis and filtering of the single-cell RNA sequencing (scRNA-seq) datasets were performed using the Seurat R package (v4.5.0)^37^. Seurat is a widely used and powerful tool for scRNA-seq analysis. We employed its functionalities for rigorous quality control, normalization, identification of sources of cellular heterogeneity, and integration across different datasets. To quantify the enrichment of network-derived modules, we calculated signature scores using the UCell algorithm^38^ (implemented via the pyucell package).Unlike methods that rely on data normalization, UCell provides rank-based scores calculated from Mann-Whitney U statistics, ensuring robustness against technical inter-sample variation. Four distinct signatures were evaluated: Betweenness and Participation coefficients for both Control and Obese conditions. To ensure cross-species consistency, the same core gene sets were applied to both human and mouse datasets, using the respective orthologous gene nomenclature.

### Weighted Gene Co-expression Network Construction and Module Detection

The construction of the gene co-expression network was performed using the Weighted Correlation Network Analysis (WGCNA) R package^39^, a statistical approach for building and analyzing transcriptomic network data. Using the filtered data as input for the WGCNA package, we first selected an appropriate soft-thresholding power (β). A power of β = 4 was chosen as it was the lowest value to achieve an approximate scale-free topology fit of R² ≈ 0.8.(Suplementary Figure 1E) This selected power was then used to transform the gene correlation matrix into an adjacency matrix. To ensure a robust calculation of gene correlations, the midweight bicorrelation method was employed^40^. This approach is less sensitive to outliers and can capture more complex relationships compared to standard correlation metrics. Following the adjacency matrix calculation, gene modules, defined as clusters of highly co-expressed genes, were identified^41^.

### Network Visualization and Functional Analysis

The resulting co-expression network and its modules were visualized using Cytoscape software (v3.9.1). Topological analysis of the network, such as connectivity and centrality metrics, was performed using the NetworkAnalyzer^42^ plugin. To infer the biological significance of the identified gene modules, functional enrichment analysis was conducted using the ClueGO^43^ plugin, which identifies and visualizes enriched Gene Ontology (GO) terms and biological pathways.

### Hub Gene Identification and selection

Hub genes within the network were identified using two complementary approaches. We first applied betweenness centrality approach from NetworkAnalyzer, a standard metric for identifying central nodes in network topology. To achieve a more nuanced classification, we also utilized the R package Rnetcarto^26^, which performs a cartographic analysis of complex networks. This method moves beyond the classical hub definition based solely on the number of connections (degree) by classifying nodes according to their participation coefficient and intra-modular connectivity^44^. This approach categorizes nodes based on their topological roles, such as their importance within their own module versus their role in connecting different modules, providing richer functional insights than centrality metrics alone.

## Results

### Defining Macrophage Networks Through Phenotypic Module Construction

Cellular responses to stimuli are not linear events but complex, interactive processes governed by intricate networks of gene expression that maintain or redefine cellular homeostasis. Understanding the principles that control these network level transitions is a central challenge in modern biology. Macrophages, with their profound functional plasticity, serve as a paradigm for this complexity, their switch to a pro inflammatory state during obesity is a key driver of metabolic disease. To dissect the molecular logic underlying this functional shift, we leveraged a systems immunology approach. By analyzing public omics data, we constructed and interrogated the gene regulatory networks to identify the core drivers of macrophage pro inflammatory polarization in obesity.

To investigate the transcriptional networks governing macrophage function in obesity, we constructed a gene co-expression network using RNA-seq data from adipose tissue macrophages isolated from mice fed a high-fat diet (HFD, n=9) or a conventional diet (ND, n=7) (GSE126407). Using Weighted Gene Co-expression Network Analysis (WGCNA), we identified 12 distinct gene modules (Supplementary Figure 2A). By correlating these modules with the obesity phenotype, we identified specific gene networks strongly associated with either the obese or lean state.

Notably, the ‘Blue’ module (MEblue) showed a near perfect positive correlation with obesity (,), while the ‘Black’ module (MEblack) also displayed a significant association (r = 0.72, p = 0.002). Conversely, the ‘Purple’ (MEpurple) and ‘Magenta’ (MEmagenta) modules were highly correlated with the lean control state (r = 0.81, p = 1×10⁻⁴ and r = 0.67, p = 0.004, respectively), hierarchical clustering of module indicated that several of these significant modules were closely related (Supplementary Figure 2B). To consolidate these findings, we merged modules with an eigengene similarity greater than 50%, resulting in six final modules (Figure 1A). Based on the strength of their phenotypic association, we selected the ‘Blue’ (obesity associated, r = 0.97, p = 5×10^−10^) and ‘Magenta’ (lean associated, r = 0.77, p = 5×10^−4^) modules for in depth analysis, positing that genes with high module membership are central to establishing the macrophage’s functional state in health and disease.

**Figure 1.**
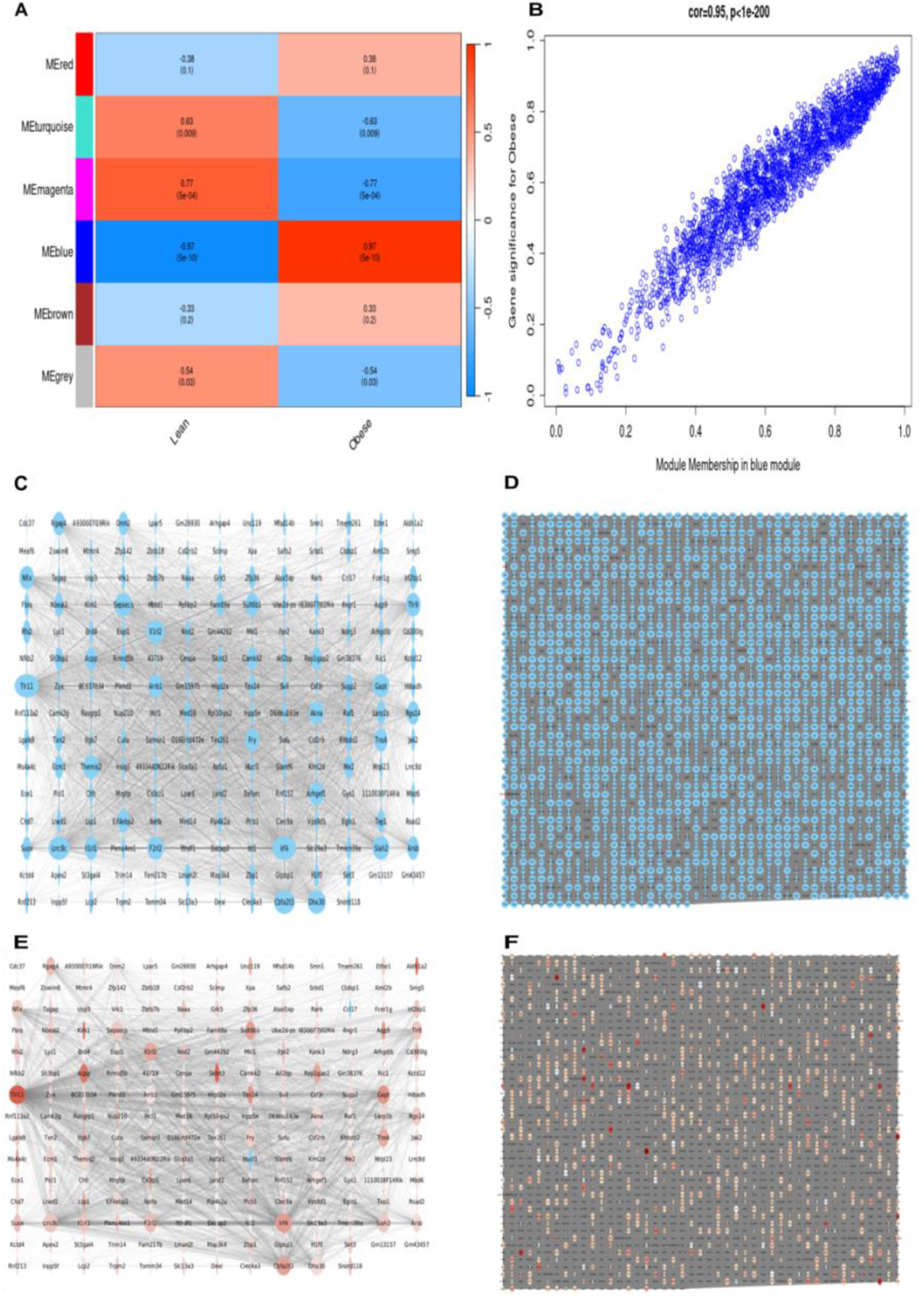
Network analysis reveals a profound remodeling of the macrophage transcriptional architecture in obesity. (A) Module-trait relationship heatmap from WGCNA of murine adipose tissue macrophage transcriptomes. The analysis identifies modules of co-expressed genes correlated with either the lean or obese phenotype. The ‘Magenta’ module was the most significant for the lean state (r = −0.77, p = 5×10⁻⁴), while the ‘Blue’ module was the most significant for the obese state (r = 0.97, p = 5×10⁻¹⁰). (B) Scatter plot showing a strong positive correlation between Gene Significance (GS) for obesity and Module Membership (MM) within the ‘Blue’ module (r = 0.95, p < 1×10⁻²⁰⁰), validating its biological relevance. (C) Co-expression network of the lean-associated ‘Magenta’ module, which is sparse and consists of 193 genes (nodes) and 3,312 connections (edges).(D) Co-expression network of the obesity-associated ‘Blue’ module, which is dramatically larger and denser, consisting of 2,091 nodes and over 1.5 million edges. (E) Visualization of differentially expressed genes (DEGs) within the ‘Magenta’ network. Red nodes indicate upregulated DEGs in the obese state. (F) Visualization of DEGs within the ‘Blue’ network, showing that while many DEGs are present, they constitute a smaller fraction of the vastly expanded network compared to the lean state.

To validate the biological relevance of these modules, we analyzed the relationship between the significance of individual genes for the clinical trait (Gene Significance, GS) and their connectivity within the module (Module Membership, MM). We found a strong positive correlation between these two metrics, confirming that genes highly connected within a module are also highly significant for the associated phenotype. Specifically, in the obesity associated ‘Blue’ module, the correlation between GS and MM was exceptionally high (r = 0.95, p < 0.001) (Figure 1B). A similar significant correlation was observed in the lean associated ‘Magenta’ module (r = 0.68, p < 0.001) (Supplementary Figure 2C). This powerful correlation demonstrates that these modules are composed of genes that are both highly co-regulated and biologically central to the disease state. Moreover, the high degree of intramodular correlation within both modules, visualized in the adjacency matrix heatmap (Supplementary Figure 2D), validates them as cohesive and robust gene sets. These modules therefore possess the ideal characteristics for network construction: strong internal correlation and a powerful, opposing association with the healthy and obese states. This robust validation prompted their selection for the in-depth construction of the core macrophage co-expression networks central to this work.

### The network architecture of the pro-inflammatory macrophage

To visualize the internal architecture of the key macrophage networks, we constructed co-expression networks from the lean associated ‘Magenta’ module and the obesity associated ‘Blue’ module using Cytoscape. This analysis revealed a profound remodeling of the macrophage transcriptional network in obesity.

The ‘Magenta’ (lean) network was relatively small and sparse, consisting of 193 nodes and 3,312 edges, with a diameter of 4 (Figure 1C). Topological analysis indicated the presence of some central, highly connected genes within this basal state (Supplementary Figure 2E). In stark contrast, the ‘Blue’ (obese) network was dramatically larger and more complex. It comprised 2,091 nodes connected by 1,575,772 edges, forming a much denser web of interactions with a network diameter of 6 (Figure 1D). The node degree and betweenness centrality distributions for the obese network were substantially shifted, revealing a large number of highly connected genes that emerge during the obese state (Supplementary Figure 2F). This extensive rewiring signifies a profound shift in the transcriptional landscape of macrophages in obesity.

We next sought to determine whether differentially expressed genes (DEGs) between the obese and lean conditions were key constituents of our identified networks. An intersectional analysis revealed that a number of DEGs were present in the 193 gene ‘Magenta’ (lean) network(Figure 1E), indicating their involvement in basal macrophage functions. A different pattern emerged in the obese state. Although the ‘Blue’ (obese) network also contained numerous upregulated DEGs (figure 1F), the overlap was less comprehensive relative to the network’s large size. This critical observation implies that the massive transcriptional reprogramming in obese macrophages is driven not only by genes that are strongly upregulated but also by a vast network of co-expressed genes that do not meet the threshold for differential expression. This suggests that co-expression, independent of large magnitude expression changes, is a key mechanism controlling the pro inflammatory macrophage profile in obesity.

Having established that co-expressed gene networks are central to macrophage identity, we sought to define their functional signatures using ClueGO pathway analysis. This revealed a clear functional polarization between the lean and obese macrophage states. The genes from ‘Magenta’ (lean) network was primarily associated with pathways of immune surveillance and transcriptional regulation, including immune receptor activity and positive regulation of the anti-inflammatory cytokine IL-10 (Figure 2A). Conversely, the vastly expanded ‘Blue’ (obese) network was overwhelmingly enriched in pathways driving aggressive inflammation and immune cell activation. Key enriched terms included the regulation of the pro inflammatory cytokine IL-6, positive regulation of leukocytes, macrophage activation, and TNF signaling(Figure 2B). This demonstrates that the transcriptional reprogramming observed in obese macrophages translates to a functional switch, redirecting the cell from a regulatory role to that of a key effector of chronic inflammation.

**Figure 2.**
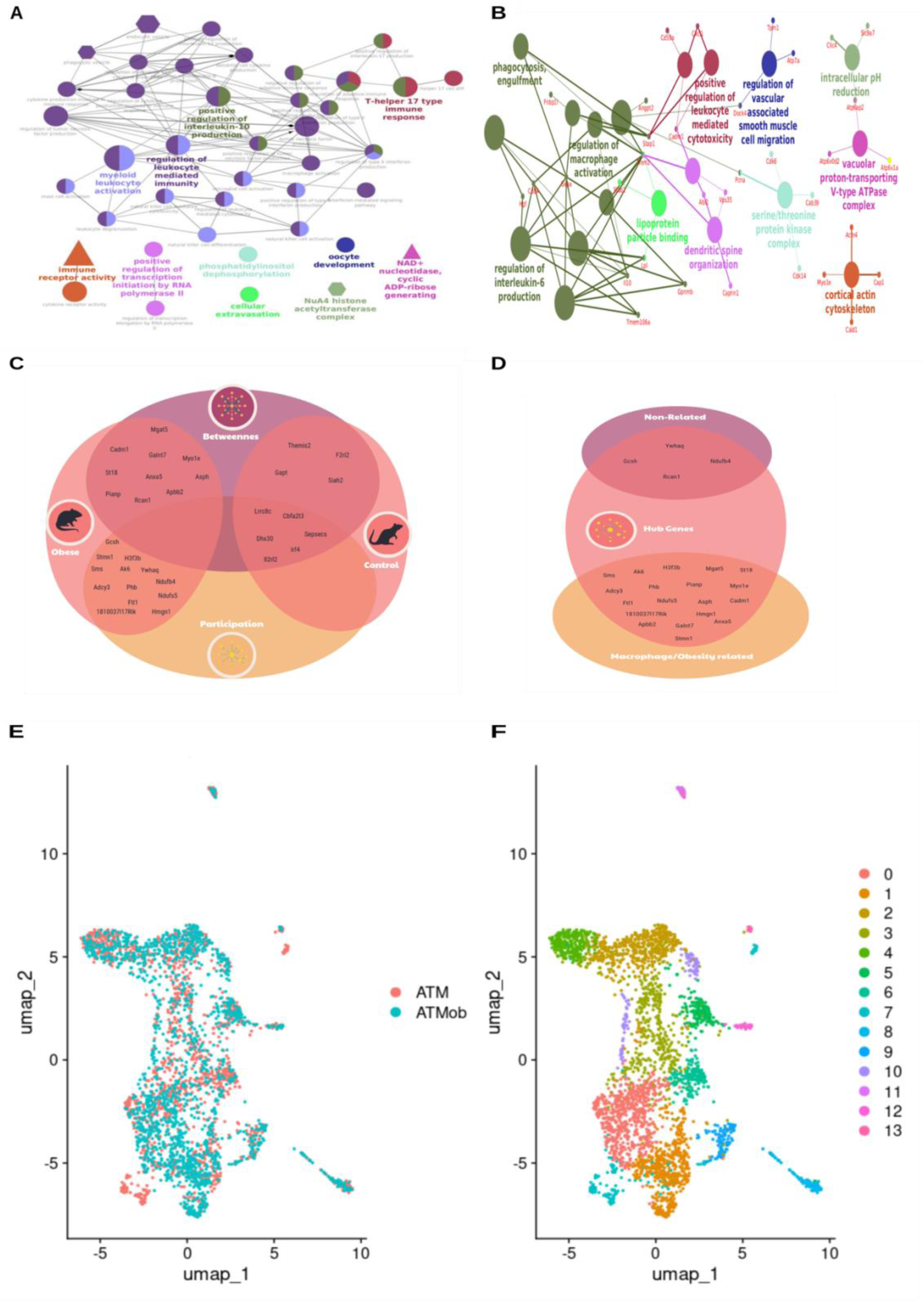
Functional polarization and hub gene identification in macrophage networks (A) Functional enrichment network (ClueGO) of the lean ‘Magenta’ module. Nodes represent enriched biological pathways, which are predominantly related to homeostatic functions like immune receptor activity and positive regulation of IL-10.(B) Functional enrichment network of the obese ‘Blue’ module, revealing a switch to a pro-inflammatory program. Enriched pathways include regulation of IL-6, macrophage activation, and TNF signaling.(C) Venn diagram illustrating the overlap and divergence of hub genes identified by two methodologies: betweenness centrality (’Betweenness’) and network cartography (’Participation’, via Rnetcarto), in both the lean (’Control’) and obese states.(D) Venn diagram showing the relationship of the identified obese hub genes to genes previously associated with macrophages or obesity in the literature. The intersection highlights the discovery of novel hub genes with no prior known link to either.(E) UMAP embedding of single-cell RNA-seq data from murine adipose tissue macrophages (ATMs), colored by sample origin: lean (ATM, teal) and obese (ATMob, red).(F) The same UMAP embedding colored by the 14 distinct subpopulations (Seurat clusters) identified by unsupervised clustering, demonstrating the cellular heterogeneity within the macrophage populations.

### Orthogonal Hub Identification Uncovers Novel Regulators of Macrophage Pathology in Obesity

To identify key regulatory genes, we first implemented a centrality based approach. In the sparsely connected lean network, this method identified 11 hub genes based on betweenness centrality (Table 1). Conversely, the densely interconnected obese network yielded over 100 nodes with high centrality. We therefore prioritized the top 12 hubs from this network based on a combined ranking of betweenness centrality and node degree (Table 1). Recognizing that centrality can overlook nodes with critical roles within network modules, we next employed a network cartography approach to classify genes based on their connectivity patterns (participation coefficient and intra-modular degree). In the lean network, this role based analysis identified seven hubs, all of which were a subset of those identified by centrality. This methodological consensus which included key immune sensors (Tlr1, Irf4) and transcriptional regulators (Dhx30, Cbfa2t3) validates the central homeostatic role of these genes.

In stark contrast, the two methods identified entirely divergent sets of hubs in the obese network. The 12 centrality based hubs were established regulators of inflammation and immune activation, such as Mgat5 (Th1 cytokine regulation), Cadm1 (T-cell activation), and Anxa5 (IL-6 signaling) (Table 1). Conversely, the 11 hubs identified by network cartography were enriched for distinct biological functions, including chromatin remodeling (H3f3b, Hmgn1), metabolic control (Ndufs5), and macrophage polarization (Sms, Phb) (Figure 2C).

Critically, our dual methodology pipeline uniquely identified four hub genes Rcan1, Ywhaq, Gcsh, and Ndufb4 with no previously established links to macrophage biology or obesity (Figure 2D). The identification of these novel regulatory nodes highlights the capacity of our integrative network analysis to uncover previously unrecognized drivers of macrophage pathology.

| Lean |  | Obese |  |
|---|---|---|---|
| Betweenness | Participation | Betweenness | Participation |
|---|---|---|---|
| Irf4 | Irf4 | Mgat5 | H3f3b |
| Tlr11 | Tlr11 | Stmn1 | Sms |
| Sepsecs | Sepsecs | Cadm1 | Ndufs5 |
| Lrrc8c | Lrrc8c | Anxa5 | Ndufb4 |
| Dhx30 | Dhx30 | Rcan1 | Ywhaq |
| Cbfa2t3 | Cbfa2t3 | Apbb2 | Gcsh |
| Gapt |  | Galnt7 | Phb |
| Siah2 |  | Asph | Ak6 |
| Il1rl2 |  | Myol1e | 1810037I17Rik |
| Themis2 |  | Pianp | Ftl1 |
| F2rl2 |  | St18 | Hmgn1 |
|  |  | Adcy3 |  |

### Single-cell resolution validates network derived hubs in pro inflammatory macrophage subpopulations

To validate and map our network hubs in situ, we analyzed a public scRNA-seq dataset (GSE117176) of adipose tissue macrophages (ATMs) from lean and obese mice. Unsupervised clustering resolved these cells into 14 distinct subpopulations, confirming the profound heterogeneity of ATMs in both metabolic states (Figure 2E-F). We then systematically interrogated this high-resolution landscape for the expression of all four candidate hub gene set those derived from both centrality and Rnetcarto analyses of the lean and obese networks (Figure 3A-B).

**Figure 3.**
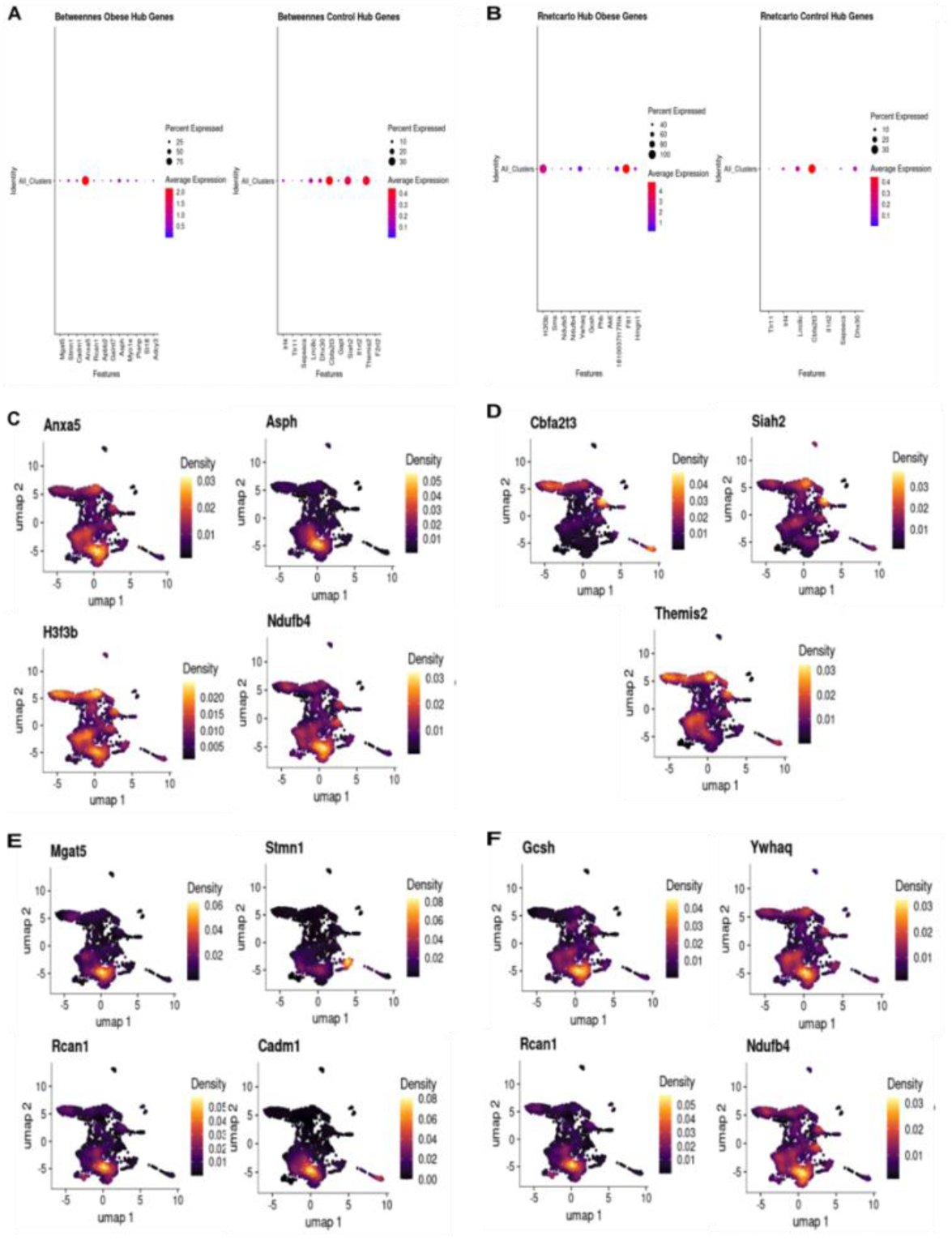
Single-cell analysis reveals distinct expression patterns of networks hubs (A) Dot plot showing the expression of centrality-derived hub genes from the obese (left) and lean (right) networks across all 14 macrophage clusters. Dot size indicates the percentage of cells expressing the gene in a cluster; color intensity indicates the average expression level.(B) Dot plot showing the expression of Rnetcarto-derived hub genes from the obese (left) and lean (right) networks across all 14 macrophage clusters.(C) UMAP plots showing the expression density of representative hubs from the obese network. Anxa5 and H3f3b show broad expression, while Asph and Ndufb4 are more restricted.(D) UMAP plots showing the expression density of key hubs from the lean network, Cbfa2t3, Siah2, and Themis2.(E) UMAP plots showing the highly concentrated, cluster-specific expression of centrality-derived obese hubs Mgat5, Stmn1, Rcan1, and Cadm1.(F) UMAP plots showing the highly concentrated, cluster-specific expression of novel Rnetcarto-derived obese hubs Gcsh, Ywhaq, Rcan1, and Ndufb4, linking them to a specific pro-inflammatory subpopulation.

This analysis revealed a clear dichotomy between hubs with specialized, cluster-specific expression and those with broader, more universal roles. In the lean state, the key homeostatic regulator Cbfa2t3 was highly concentrated in a specific subpopulation, whereas hubs like Siah2 and Themis2 were more widely distributed. Other lean network hubs, including Il1rl2 and Tlr11, were also restricted to discrete clusters, suggesting specialized functions (Figure 3D, Supplementary Figure 3A).

This dichotomy was even more pronounced in the obese state. Most centrality derived hubs, including Mgat5, Stmn1, and the novel hub Rcan1, exhibited highly concentrated expression, pinpointing them to a specific pro-inflammatory macrophage subpopulation (Figure 3E). In stark contrast, the hub gene Anxa5 was broadly expressed across most cells, suggesting a more general role in the inflammatory response (Figure 3C).

The Rnetcarto derived hubs further clarified this pattern. A subset, including H3f3b and Ftl1, were expressed broadly, likely representing core cellular machinery co-opted during inflammation. However, a distinct group including metabolic regulators like Ndufs5 co-localized within the same specialized macrophage cluster (Supplementary Figure 3B). Critically, our novel hub genes Ywhaq, Gcsh, and Ndufb4 also displayed this highly concentrated expression pattern, definitively linking them to this specific, pathogenic macrophage subpopulation that emerges in obesity (Figure 3F).

Together, these high resolution data validate our network-derived hubs by assigning them to discrete macrophage subpopulations with distinct, inferred functional roles distinguishing specialized drivers of pathology from genes with broader functions in the tissue microenvironment.

### Translational validation of network hubs in human adipose tissue macrophages

To determine the clinical relevance of our findings, we next investigated the expression of the murine hub genes in human adipose tissue macrophages (hATMs). We identified human orthologs for our candidate genes and analyzed their expression in a public scRNA-seq dataset (GSE176171) derived from human adipose tissue.(Supplementary Figure 4A).Macrophages from this dataset were stratified into lean and obese groups based on patient Body Mass Index (BMI), resulting in 6,285 lean and 11,027 obese hATMs for analysis (Figure 4A). We then assessed the expression of the Rnetcarto derived hub gene orthologs within this human cohort. Despite some murine genes lacking human orthologs or being absent from the dataset, key findings from the mouse network were conserved.

**Figure 4.**
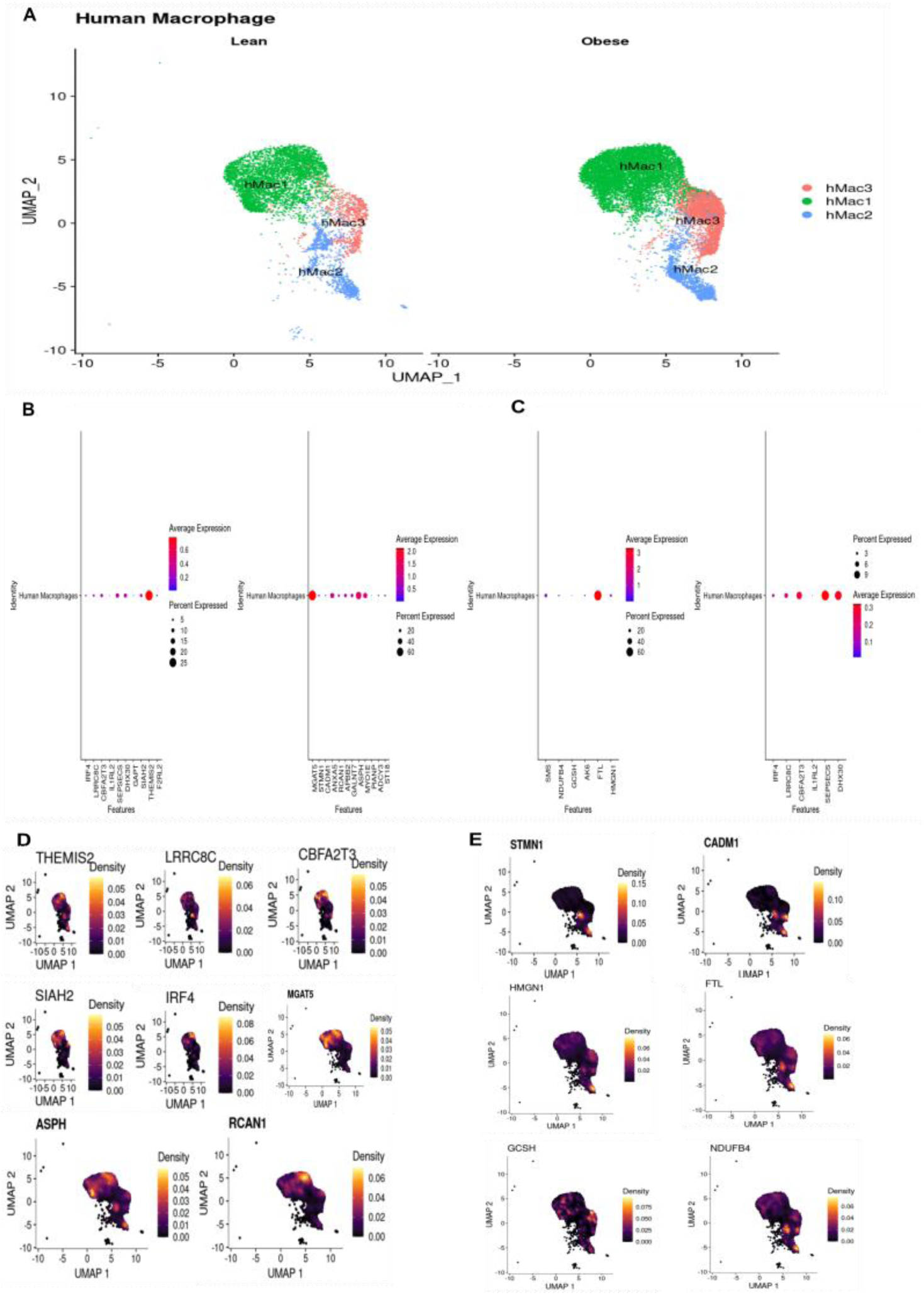
Single-cell analysis of human macrophages validates the conservation of network hubs (A) UMAP embeddings of human adipose tissue macrophages (hATMs) from a public scRNA-seq dataset (GSE176176), stratified by patient BMI into lean (left; 6,285 cells) and obese (right; 11,027 cells) groups. Cells are colored by macrophage sub-cluster identity (hMac1, hMac2, hMac3).(B) Dot plots showing the expression of human orthologs of the centrality-derived murine hubs in human macrophages. Obese-network hubs are on the left; lean-network hubs are on the right.(C) Dot plots showing the expression of human orthologs of the Rnetcarto-derived murine hubs in human macrophages. Obese-network hubs are on the left; lean-network hubs are on the right.(D) UMAP plots showing the expression density of representative human hub orthologs. Hubs defining the lean state, like THEMIS2 and CBFA2T3, show broad expression, while others like SIAH2 and IRF4 are more restricted. Key obese hubs like MGAT5, RCAN1, and ASPH are also validated.(E) UMAP plots showing the highly concentrated, cluster-specific expression of human orthologs for key obese-network hubs, including STMN1, CADM1, HMGN1, FTL, and the novel hubs GCSH and NDUFB4.

This cross-species analysis revealed a striking conservation of the key regulatory hubs. Our analysis confirmed that the key transcriptional signatures are conserved. The proinflammatory hubs from the obese murine network were prominently expressed in macrophages from obese humans, including MGAT5, ASPH, ANXA5, and the Rnetcarto hub FTL1. Importantly, our novel hub RCAN1 was also validated in this obese human context (figure 4B). Similarly, the homeostatic hubs from the lean murine network were characteristic of human lean macrophages, with THEMIS2 and CBFA2T3 being notable examples. Two of our other novel hubs, GCSH and NDUFB4, were also present in the human dataset, though YWHAQ was not detected.(Figure 4C) This cross-species validation demonstrates that the regulatory hubs identified by our network approach, including several previously uncharacterized genes, are fundamental to macrophage function in human obesity.

### Human single-cell analysis reveals distinct expression patterns of conserved hubs

To validate the human relevance of our hubs, we next examined the expression density and distribution of their orthologs in the human macrophage scRNA-seq data. This analysis revealed that the hubs defining the lean and obese states exhibit distinct expression patterns, suggesting different functional roles.

In human lean macrophages, the homeostatic signature was defined by two classes of hubs. A core group, including THEMIS2, LRRC8C, and the key regulator CBFA2T3, were broadly expressed across most cells, likely representing a conserved, global regulatory program. In contrast, other hubs like SIAH2 and IRF4 showed expression restricted to discrete subpopulations, suggesting specialized functions even within the healthy state. The hubs associated with obesity also segregated into two distinct functional groups. A subset of centrality derived hubs, including MGAT5, ASPH, and our novel hub RCAN1, were broadly expressed across the obese macrophage landscape, indicative of a widespread pro inflammatory program(Figure 4D). However, a second, larger group of hubs from both centrality and Rnetcarto analyses showed highly concentrated expression, co-localizing to specific, distinct subpopulations. This specialized group includes established inflammatory drivers like STMN1 and CADM1, as well as Rnetcarto-derived hubs HMGN1 and FTL(figure 4E).

Critically, our novel hub genes validated in humans, GCSH and NDUFB4, also displayed this highly restricted expression pattern, definitively linking them to these specialized, likely pathogenic, macrophage subpopulations that expand in human obesity(Figure 4 E). This dichotomy between broadly expressed and cluster-specific hubs reveals a complex regulatory architecture, where a general inflammatory state is likely driven and sustained by highly specialized macrophage subsets defined by a unique cohort of regulatory genes. Another genes are showed in Suplementary Figure 4B-D.

### Hub-set signature scores map the lean and obese network programs onto discrete macrophage subpopulations

To test whether the hubs behave as coordinated programs rather than isolated genes, we scored every cell with four rank-based UCell signatures built from Table 1: lean and obese hubs, each defined by betweenness or participation. In murine ATMs, both lean signatures were low in most cells and confined to a few groups, mainly the upper region of the UMAP and small satellite clusters (Fig. 5A,C;). In contrast, both obese signatures were raised across the macrophage landscape (Fig. 5B,D), peaking in [hMac3], the same subpopulation where Mgat5, Stmn1, Rcan1 and Ndufb4 were concentrated (Fig. 3E-F). The obese participation score had a high baseline in nearly all cells, which suggests that broadly expressed hubs (e.g., H3f3b, Ftl1) contribute to it, with more restricted hubs adding intensity in specific cells.

**Figure 5.**
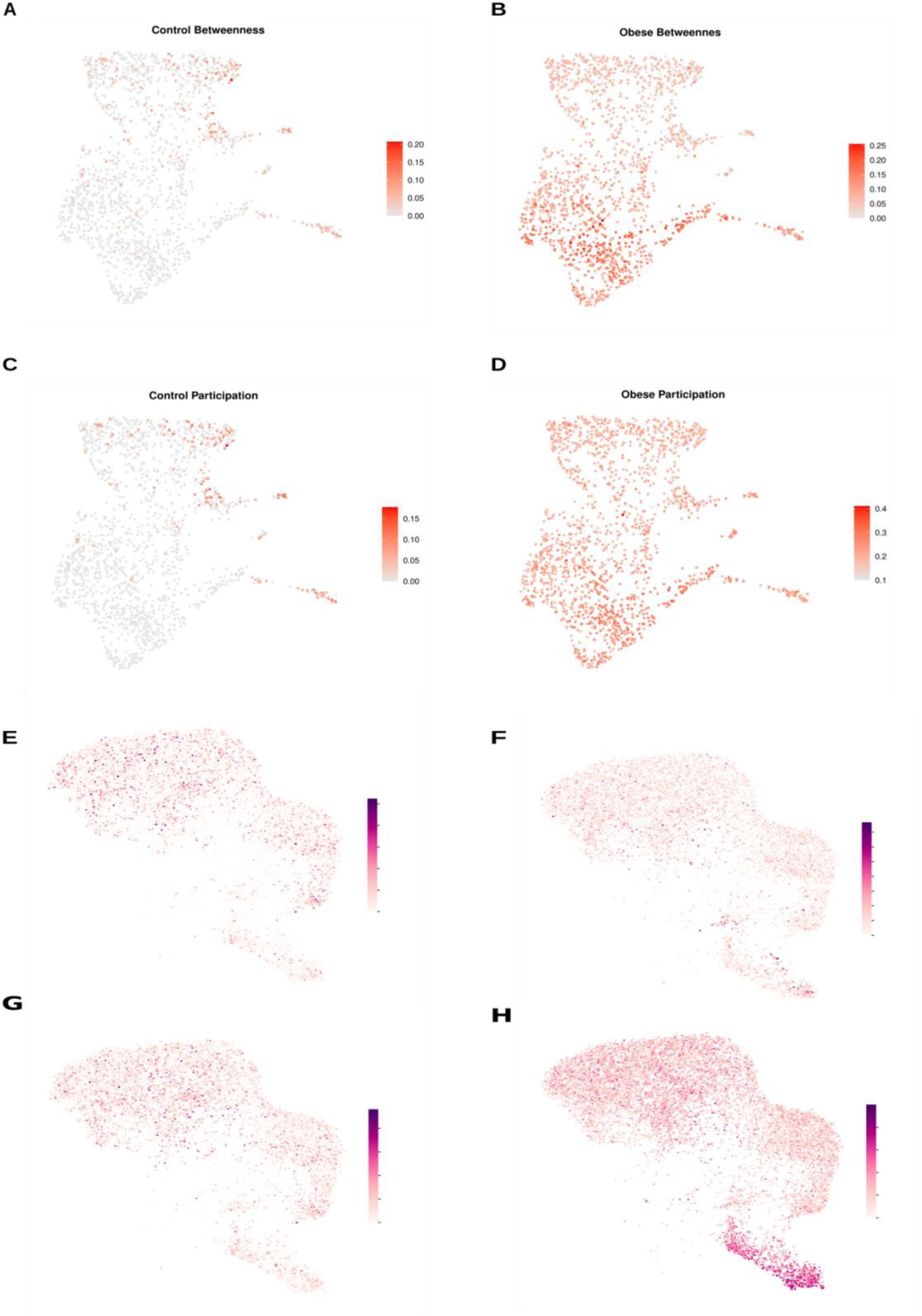
Hub-set signature scores in murine and human adipose tissue macrophages. UCell scores per cell for the lean (control) and obese hub sets in Table 1. (A-D) Murine ATMs (GSE117176): lean betweenness (A), obese betweenness (B), lean participation (C), obese participation (D). (E-H) Human ATMs (GSE176171), same order. Each dot is a cell; colour is the UCell score participation hubs.(H) UMAP embeddings of human adipose tissue macrophages (hATMs) enriched from obese participation hubs.

In human ATMs, the lean signatures were low and scattered (Fig. 5E,G) and the obese betweenness signature was weak and diffuse (Fig. 5F). The obese participation signature (Fig. 5H) was higher overall and formed a sharp hotspot in a discrete [hMac cluster; % of cells; statistics]. Because the UMAP spaces of the two species are independent, we did not infer correspondence from position;

## Discussion

Obesity has become a global health crisis, fueling a spectrum of chronic inflammatory diseases^45–50^, yet the genetic and molecular drivers that link excess adiposity to systemic pathology are not fully resolved. A central event in this process is the accumulation and proinflammatory activation of macrophages within adipose tissue ^4,51^. However, a significant gap has persisted in our understanding of the gene networks that orchestrate this cellular reprogramming. Here, we bridged this gap by combining transcriptome-wide network analysis with high resolution single cell validation. This dual approach allowed us to move beyond individual gene effects and identify the core regulatory modules and previously uncharacterized hub genes that define the proinflammatory macrophage identity in obesity.

While co-expression network analysis has been applied to fields like oncology^52^, its application to macrophage biology in obesity remains nascent. A key advance of this study is the identification of previously uncharacterized genes Rcan1, Ywhaq, Gcsh, and Ndufb4 as central regulatory hubs in adipose tissue macrophages. While our network analysis also confirmed the importance of genes with established roles in immunity or obesity, such as Stmn1, Asph, and Hmgn1, the discovery of these novel players opens new avenues for understanding the molecular basis of chronic inflammation. These insights were enabled by a dual methodology approach that provided a more comprehensive view of network architecture than is possible with standard methods alone. Classical centrality metrics successfully identified key homeostatic regulators (Cbfa2t3, Siah2) and established pro inflammatory drivers (Stmn1, Cadm1, Anxa5). However, the role based cartographic analysis (Rnetcarto) was uniquely able to uncover a different class of hubs, including epigenetic regulators (H3f3b, Hmgn1), metabolic genes, and several of the novel candidates (Ywhaq, Gcsh). This demonstrates that combining these complementary approaches is a powerful strategy to deconstruct complex biological networks, revealing distinct layers of regulation that would otherwise be missed.

The hubs defining the homeostatic lean macrophage network are pleiotropic regulators with established roles in both maintaining physiological balance and driving pathology when dysregulated. The consensus between our two network identification methods highlights the central importance of these genes. For instance, the transcriptional co-repressor Cbfa2t3, which we identified as a key hub, is not only critical for development^53,54^ but has also been implicated in leukemia ^55^ and, conversely, in promoting healthy metabolic phenotypes during weight loss^56^.

This theme of dual functionality extends to other hubs. Siah2 and Themis2, for example, are critical for inflammation and immune cell activation in contexts ranging from adipose tissue to cancer^57–60^. Similarly, genes involved in core cellular processes, such as transmembrane transport (Lrrc8c) and RNA processing (Dhx30), also have context-dependent roles in regulating inflammatory signals^61–63^. This suggests that the homeostatic state of lean macrophages is actively maintained by a network of powerful, multi functional hubs whose balanced expression is critical for preventing the transition to a pathological state. The centrality derived hubs of the obese macrophage network constitute a molecular toolkit for orchestrating tissue inflammation and metabolic dysfunction. This network integrates signals for immune cell activation and adhesion, driven by hubs such as Stmn1 and Cadm1^64,65^, with the machinery for cellular effector functions like phagocytosis, governed by Myo1e^66,67^.

Crucially, this inflammatory program is interwoven with direct drivers of metabolic disease. We identified Asph, a gene linked to M1 macrophage polarization^68,69^, and the calcineurin regulator Rcan1, a known player in type 2 diabetes^70,71^. The identification of Rcan1 is particularly significant given recent evidence linking it to lipid accumulation^72^, positioning it as a key node connecting inflammation to the metabolic consequences of obesity. The convergence of these diverse pathological functions onto a small set of hub genes highlights a coordinated transcriptional program that reinforces the pro inflammatory, metabolically disruptive phenotype of macrophages in obese adipose tissue. Our role-based network analysis using Rnetcarto uncovered the hubs that form the epigenetic and metabolic core of the pathological obese macrophage. This regulatory layer is anchored by chromatin-modifying proteins like H3f3b and Hmgn1, the latter of which also functions as a potent alarmin linking nuclear state to innate immune activation^73,74^. This epigenetic reprogramming is tightly integrated with a metabolic rewiring centered on mitochondrial hub Ndufs5, which are tied to obesity related comorbidities like diabetes and atherosclerosis ^75^,Ndufb4 is a nuclear gene encoding a subunit of the mitochondrial NADH:ubiquinone oxidoreductase (complex I). In murine models, it has been genetically linked to susceptibility to diabetes^76^. Conversely, in humans, it has been identified as a potential biomarker associated with hypertension77. Intriguingly, despite the strong epidemiological links between diabetes and hypertension, a direct association between Ndufb4 and obesity or macrophage driven inflammation has not been established. This pattern of co-expression across different but related metabolic pathologies positions Ndufb4 as a potential hub gene within a network of interconnected metabolic diseases, warranting further investigation into its role in obesity. The hubs identified by Rnetcarto reveal the deep seated reprogramming of the obese macrophage, linking its metabolic state directly to its effector functions. We identified novel hubs, such as Ywhaq and Gcsh, that have established roles in systemic insulin signaling and metabolism^78,79^, suggesting they act as key sensors that translate metabolic stress into a macrophage specific response. This response is then shaped by functional hubs like Ftl1 and 1810037I17Rik, which are known to modulate M1/M2 polarization and calcium signaling^80–82^.

This entire program appears to be coordinated by master regulators like Hmgn1, a pleiotropic hub that bridges epigenetics and immunity. Hmgn1 not only modifies chromatin structure^83^ but also acts as a potent alarmin to drive TLR4-mediated inflammation^73,74^. Given that its deletion protects against obesity and insulin resistance^84,85^, its identification as a top hub in our network provides a direct mechanistic link between the nuclear, metabolic, and inflammatory dysregulation that defines the pathological macrophage in obesity.

Scoring the hubs as modules adds a second line of evidence to the network analysis. The obese hub sets in both species, whereas lean hubs marked a smaller fraction of cells, consistent with the expansion from 193 to 2,091 nodes (Fig. 1). The two hub definitions behaved differently. This matters for the novel genes, because Rcan1 belongs to the betweenness set and Ndufb4, Ywhaq and Gcsh to the participation set, and the human participation hotspot (Fig. 5H) is where [interpretation]. A set score does not show which gene drives it, so these patterns are associative and need per-gene and perturbation tests.

In conclusion, our study provides a systems level blueprint for the transcriptional reprogramming of macrophages in obesity. We demonstrate that the shift to a proinflammatory state is not a stochastic process but is orchestrated by a robust co-expression network governed by a cohort of regulatory hub genes. The conservation of key hubs from homeostatic regulators like THEMIS2 to proinflammatory drivers like MGAT5 and FTL1 in human single cell data underscores the clinical relevance of this network architecture. We propose that this cohort of hubs, including established players like Hmgn1 and the novel regulators we identified such as Rcan1, Gcsh, Ywhaq, and Ndufb4, acts as a coordinating committee that collectively enforces the chronic, low-grade inflammation characteristic of obesity. These hubs therefore represent a rich set of previously unrecognized therapeutic targets for selectively reprogramming macrophage function to ameliorate metabolic disease.

## Acknowledgements

This study was supported by Conselho Nacional de Desenvolvimento Científico e Tecnológico (CNPq) and Fundação de Amparo à Pesquisa do Estado de Minas Gerais (FAPEMIG).

## Competing interests

The authors declare no competing interests.

## Notes

https://www.ncbi.nlm.nih.gov/geo/query/acc.cgi?acc=GSE126407

https://www.ncbi.nlm.nih.gov/geo/query/acc.cgi?acc=GSE176171

https://www.ncbi.nlm.nih.gov/geo/query/acc.cgi?acc=GSE117176

### Competing Interest Statement

The authors have declared no competing interest.

