## Supplementary Materials for "Systems Immunology in Obesity: Uncovering Hub Gene Networks in Macrophage-Specific Drivers of Chronic Inflammation"

### Supplementary Figure 1

*Supplementary Figure 1 Quality control and parameter selections for WGCNA (A) Hierarchical clustering dendrogram of the initial 17 macrophage samples based on their transcriptomic profiles.(B) The same dendrogram as in (A) showing the cutoff threshold (red line) used to identify outliers. The sample VAM.2.ND.1 was identified as an outlier and removed from subsequent analysis. (C) Hierarchical clustering dendrogram of the remaining 16 high-quality samples after outlier removal. (D) Sample dendrogram and trait heatmap. The clustering of the final 16 samples is shown in relation to the clinical trait (obesity), demonstrating a clear separation between lean (white) and obese (red) samples.(E) Analysis for the selection of the soft-thresholding power (β). The left panel shows the scale-free topology fit index ($R^2$) as a function of the soft-thresholding power. The right panel shows the mean connectivity as a function of the soft-thresholding power. A power of β = 4 was selected as it is the lowest power to achieve an $R^2$ value above 0.8 while maintaining adequate connectivity.*

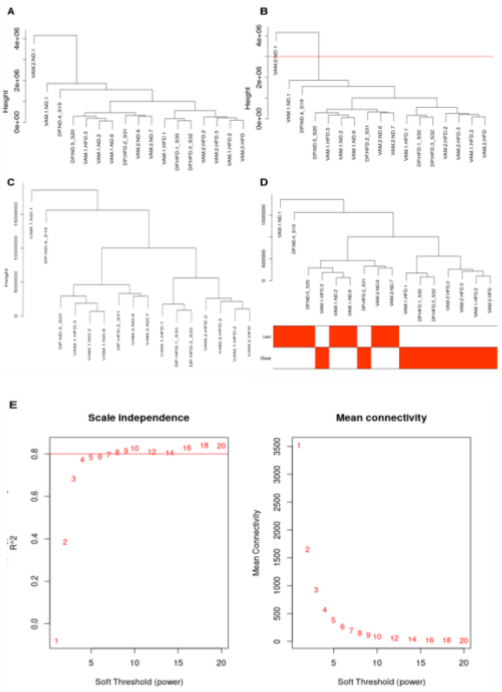


### Supplementary Figure 2

*Supplementary Figure 2 WGCNA module validation and network topology (A) Module-trait relationship heatmap showing the correlation of the initial 12 identified gene modules with the lean and obese phenotypes.(B) Hierarchical clustering dendrogram of the 12 module eigengenes. This analysis was used to* *identify and merge modules with high similarity (eigengene similarity > 50%).(C) Scatter plot showing the significant positive correlation between Gene Significance (GS) for the lean state and Module Membership (MM) within the 'Magenta' module ($r = 0.68$, $p = 2.1 \times 10^{-30}$), validating its biological relevance.(D) Heatmap of the module adjacency matrix, including the clinical trait of body weight. This plot demonstrates the high intramodular correlation within each module and confirms the strong positive and negative associations of the key modules with body weight. (E) Scatter plot showing the relationship between betweenness centrality and edge count (degree) for the 193 nodes in the lean 'Magenta' network.(F) Scatter plot showing the relationship between betweenness centrality and edge count for the 2,091 nodes in the obese 'Blue' network, illustrating the substantial increase in highly connected, central nodes.*

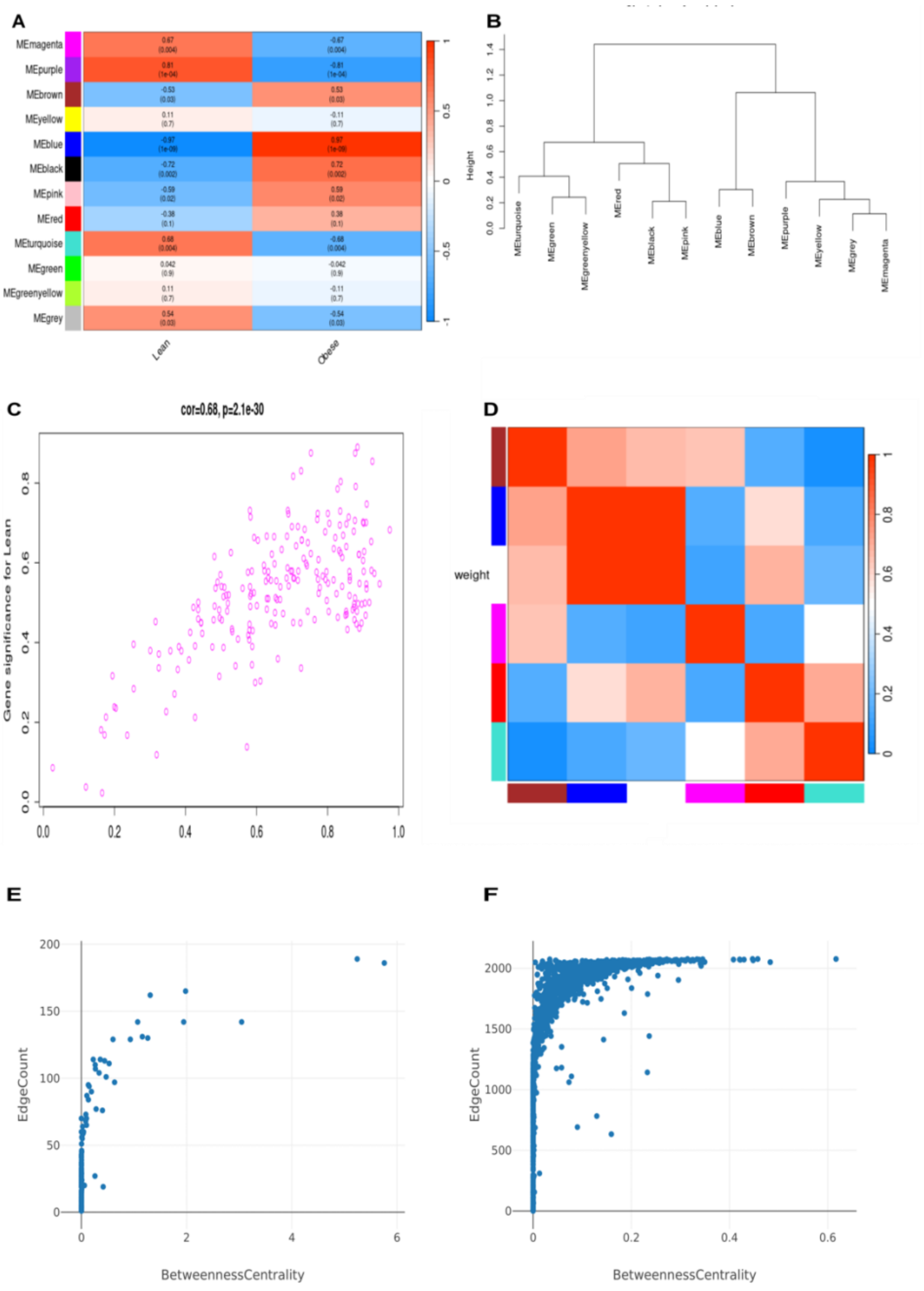


Supplementary Figure 3

*Supplementary Figure 3 High-resolution expression mapping of additional network hub genes. (A) UMAP plots showing the expression density of hub genes identified in the lean ('Magenta') network. Genes such as Tlr11 and Il1rl2 show expression restricted to discrete subpopulations, while others like Cbfa2t3 and Themis2 are more broadly distributed across the macrophage landscape.(B) UMAP plots showing the expression density of hub genes identified in the obese ('Blue') network by the Rnetcarto method. This set includes broadly expressed genes (H3f3b, Ftl1) and genes with highly concentrated expression in specific clusters, including the metabolic regulators (Ndufs5, Ndufb4) and the novel hubs (Ywhaq, Gcsh).*

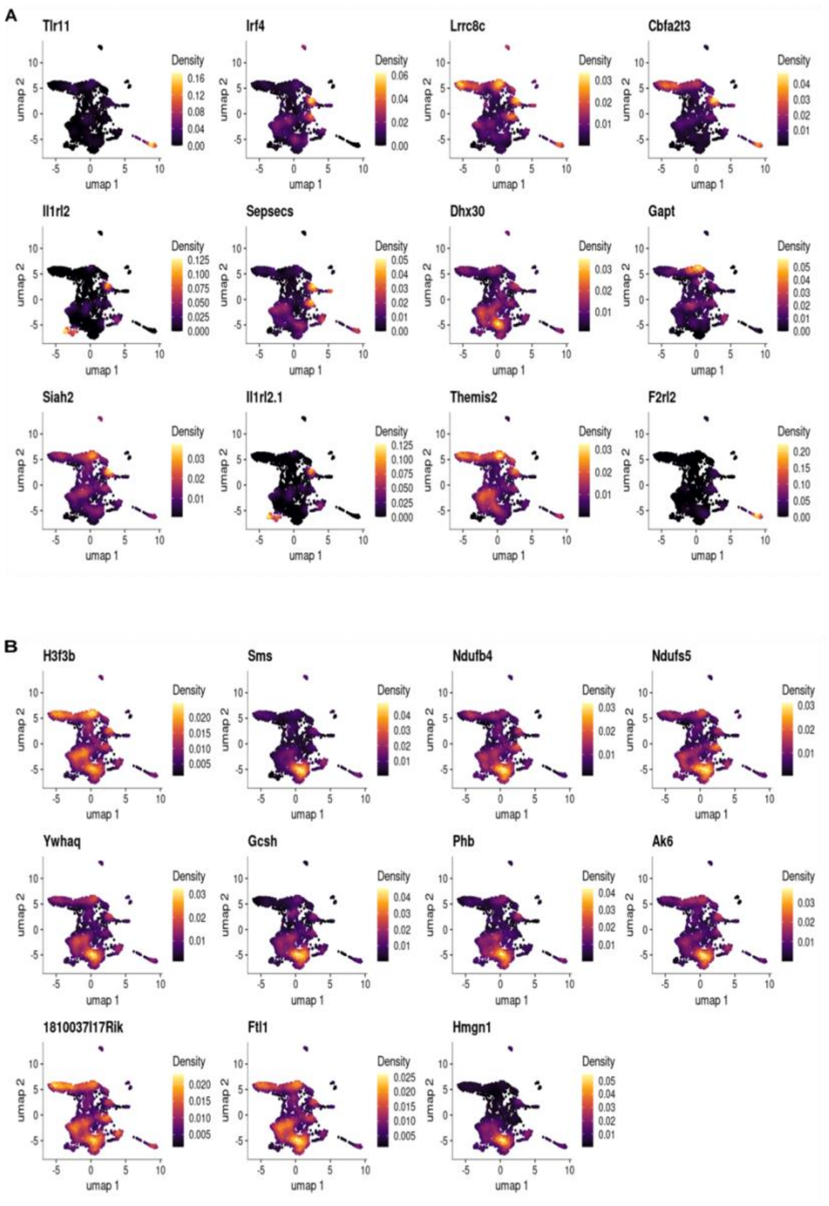


Supplementary Figure 4

*Supplementary Figure 4 Single-cell expression of human hub gene orthologs in adipose tissue*

*(A) UMAP embedding of all immune and stromal cells from the human adipose tissue scRNA-seq dataset (GSE176171). The macrophage clusters (hMac1, hMac2, hMac3) used for the main analysis are highlighted.(B) UMAP plots showing the expression density of human orthologs for the hub genes identified in the murine lean ('Magenta') network.(C) UMAP plots showing the expression density of human orthologs for the hub genes identified in the murine obese*

*('Blue') network by betweenness centrality.(D) UMAP plots showing the expression density of human orthologs for the hub genes identified in the murine obese ('Blue') network by the Rnetcarto method.e*

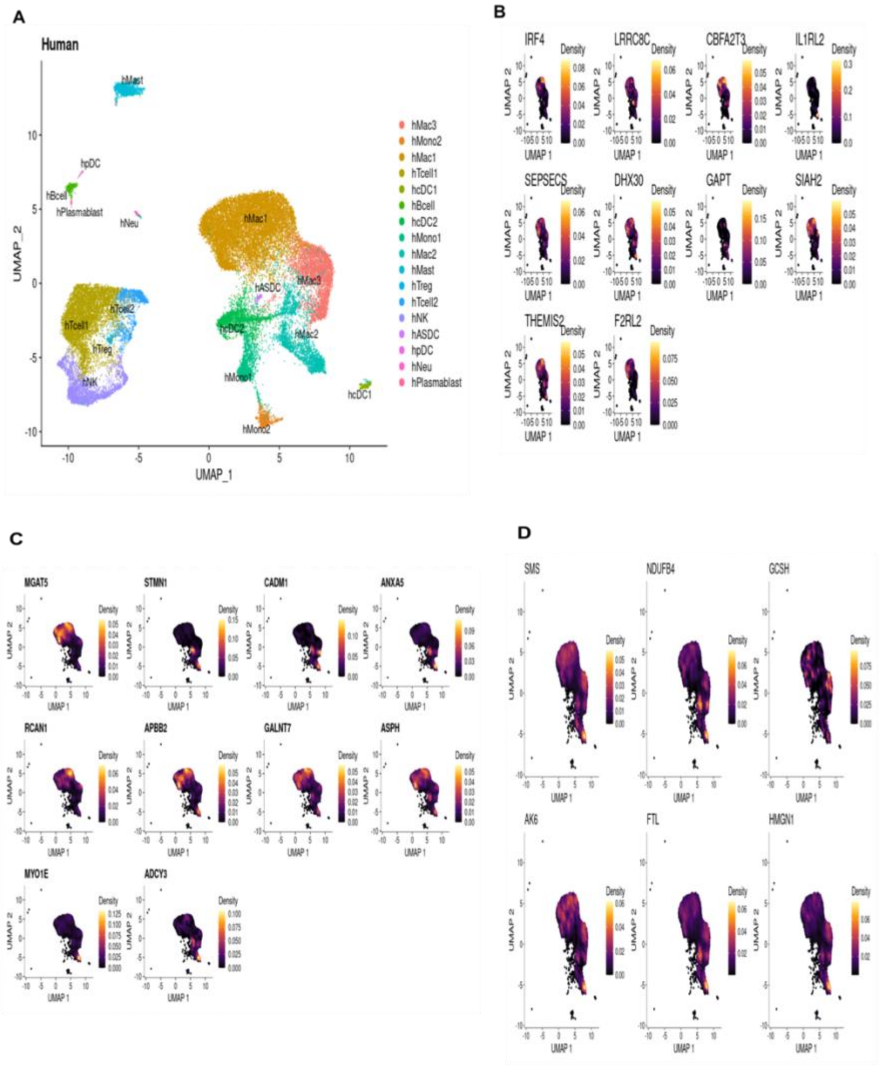
